# Sub-lethal pyrethroid exposure and ageing lead to pronounced changes in gene expression in insecticide resistance *Anopheles coluzzii*

**DOI:** 10.1101/2020.08.14.250852

**Authors:** V A Ingham, F Brown, H Ranson

## Abstract

**Background:** Malaria control is heavily reliant on the use of insecticides that target and kill the adult female Anopheline vector. The intensive use of insecticides of the pyrethroid class has led to widespread resistance in mosquito populations. The intensity of pyrethroid resistance in some settings in Africa means mosquitoes can contact bednets treated with this insecticide class multiple times with minimal mortality effects. However, the sublethal exposure may affect other traits controlling the vectorial capacity or fitness of the mosquito.

**Results:** Here we show that sublethal exposure of a highly resistant *Anopheles coluzzii* population originally from Burkina Faso to the pyrethroid deltamethrin results in large and sustained changes to transcript expression. We identify five clear patterns in the data showing changes to transcripts relating to: DNA repair, respiration, translation, ion transport and oxioreductase processes. Further, we highlight differential regulation of transcripts from detoxification families previously linked with insecticide resistance, in addition to clear down-regulation of the oxidative phosphorylation pathway both indicative of changes in metabolism post-exposure. Finally, we show that both ageing and diel cycle have major effects on known insecticide resistance related transcripts.

**Conclusion:** Sub-lethal pyrethroid exposure, ageing and the diel cycle results in large-scale changes in the transcriptome of the major malaria vector *Anopheles coluzzii*. Our data strongly supports further phenotypic studies on how transcriptional changes such as reduced expression of the oxidative phosphorylation pathway or pyrethroid induced changes to redox state might impact key mosquito traits, such as vectorial capacity and life history traits.

## Background

Insecticide based vector control tools are the cornerstone of malaria control programmes and have proven to be the most efficient means for reducing malaria related morbidity and mortality since the turn of the century [1]. However, following dramatic reductions in malaria cases since 2000, progress has plateaued in the last two years [2]; a key driver of this is widespread insecticide resistance in Anopheline vectors [3–5]. Over 2 billion insecticide treated bed nets (ITNs) have been distributed in Africa, the WHO region accounting for the majority of the malaria burden worldwide; these nets are all treated with the pyrethroid class of insecticide. Resistance to pyrethroids is ubiquitous across sub-Saharan Africa. Indeed, of the reporting countries, almost 90% detailed pyrethroid resistance [2]. In some regions, the strength of this resistance allows mosquitoes to survive multiple bed net exposures with no observable impact on mosquito longevity [6]. Pyrethroid resistance reduces the personal protection provided by bed nets but also importantly erodes the community protection afforded to non-net users by insecticide induced mortality, which has been critical for their success [7–9]. To address this problem, net manufacturers have developed new classes of nets, several of which have already been pre-qualified by WHO and are now in use in Africa. Critically, these ITNs all still contain pyrethroid insecticides but their efficacy against pyrethroid resistant mosquitoes is enhanced by the presence of a second chemistry, either an insecticide, synergist or insect sterilising agent [5, 10]. Hence, pyrethroids will remain an essential critical chemistry for malaria prevention for the foreseeable future and thus understanding the effects of pyrethroid exposure and pyrethroid resistance on *Anopheles* mosquitoes is of fundamental importance.

Pyrethroid resistance is multifactorial and is presently thought to be driven by four mechanisms; mutations to the target site of the pyrethroid insecticide, known as knockdown resistance (kdr) [11]; changes to the thickness of the mosquito cuticle that reduce penetrance of the insecticide [12]; sequestration by chemosensory proteins (CSPs) in the legs [13]; and finally, increased metabolic breakdown and clearance of the insecticide through over-expression of detoxification gene families [14–16]. Several members of the *Anopheles* cytochrome p450 family (P450s) have been shown to directly metabolise pyrethroids [16]; other detoxification gene families have also been implicated in resistance including glutathione-s-transferases (GSTs) [17], ABC transporters (ABCs) [18], carboxylesterases (COEs) [19] and UDP-glucuronyl transferases (UGTs) [20]. All these mechanisms, with the exception of *kdr*, are caused by over-expression of specific members of these gene families within resistant mosquitoes and have been identified in multiple transcriptomic datasets comparing resistant and susceptible populations [21]. The large library of transcriptomic datasets available comparing resistant and susceptible mosquitoes represents a valuable resource for identifying resistance associated genes. However, these experiments were designed to remove potential confounding induction effects of pyrethroid exposure and in most cases mosquitoes were harvested for RNA extraction 48 hours after exposure [22]. The process of correcting for induction effects loses data about how insecticide exposure could potentially affect mosquito biology and behaviour within this window. These facets of the mosquito response are important to investigate both to understand the mechanisms underpinning any post exposure behavioural changes such as willingness to blood feed [23] and to predict potential impacts of insecticide exposure on the development of the malaria parasite in the mosquito.

Previous studies have looked at the induction effects of insecticides on specific genes of interest and shown that both constitutive overexpression and induction are important in response to insecticide exposure. These studies include pyrethroid induction of cytochrome p450s in *Cx. quinquefasciatus* [24, 25] and *D. melanogaster* [26], ABC transporters in *An. stephensi* [27], CSPs in *An. gambiae* [13], COEs in *Musca domestica* [19], UGTs in *Spodeoptera exigua* [28] and GSTs in *Bactrocera dorsalis* [29]. Many of these insecticide-induced changes in transcript expression are linked with oxidative stress and the *cnc-Nrf1* pathway, which has been shown to be constitutively up-regulated in insecticide resistance *An. gambiae* and *D. melanogaster* [30–32]. As far as we are aware, no studies have looked at overall change in the whole transcriptome over an extended time course; this is important to understand the molecular response to sub-lethal insecticide exposure.

In this study we exposed 3-day old *An. coluzzii* females from a highly resistant colony established from Burkina Faso [33] to the pyrethroid insecticide deltamethrin and investigated changes in the transcriptome over a 72 hour time course. We identified five stages to the pyrethroid response, including a sustained change in genes associated with respiratory function transcription, DNA damage and translation. The experimental design also captures the effects of both ageing and diel cycle and reveals multiple genes previously associated with insecticide resistance are differentially expressed following pyrethroid exposure, ageing and throughout the diurnal cycle.

## Results

The experiments were designed to test three separate hypotheses: (i) Pyrethroid exposure induces changes to transcript expression over time; (ii) Ageing increases susceptibility to insecticides due to changes in expression of insecticide related transcripts and (iii) Diel cycle controls the expression of insecticide resistance transcripts. All experiments used the pyrethroid resistant VK7 strain of *Anopheles coluzzii*, originally colonised from Burkina Faso [33].

### (i) Pyrethroid exposure induces changes to transcript expression over time

To identify changes in transcript expression associated with pyrethroid exposure, 3-day old females were exposed to 0.05% deltamethrin papers for one hour using a standard WHO tube assay. Mosquitoes were then harvested for RNA extraction at 10 time points post exposure: 0 minutes, 30 minutes, 1-hour, 2-hours, 4-hours, 8-hours, 12-hours, 24-hours, 48-hours and 72-hours (Figure 1). A total of 8832 transcripts (9316 probes) showed differential expression compared to an unexposed control (taken before exposure) in at least one of these time points. Two separate analyses were then used. Firstly, significance-independent soft clustering of these transcripts by temporal changes in expression was performed using Mfuzz with 20 clusters (Additional File 1; Additional File 2). Exploration of these clusters using enrichment analyses identified five key trends within the dataset (Figure 1; Additional File 2). Secondly, transcripts showing significant differential expression following the same directionality over multiple time points were extracted; these transcripts are likely to represent the most important in the response to insecticide exposure.

**Figure 1:**
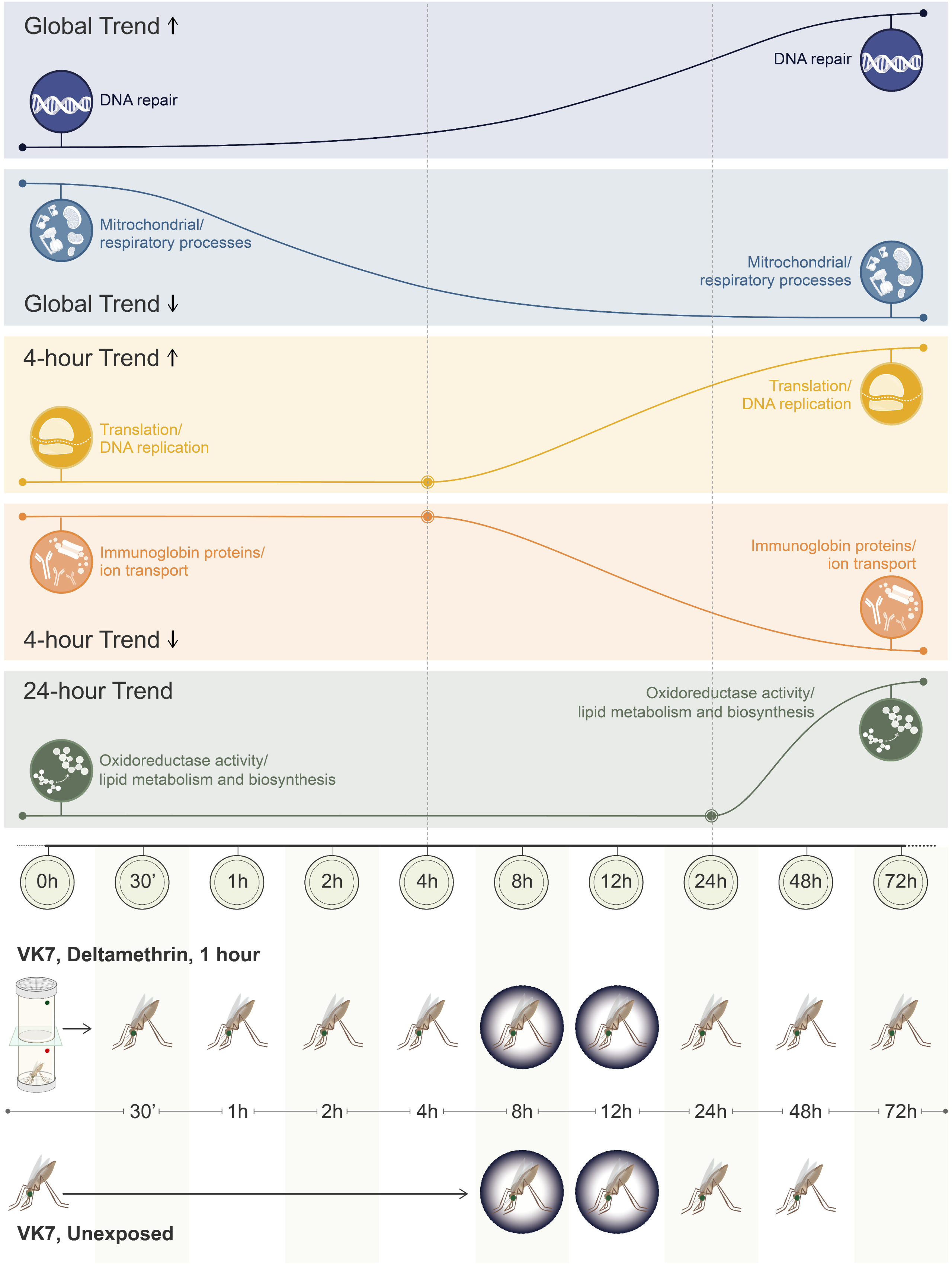
Time series trends. 5 rows demonstrating the temporal transcript pattern change and the associated enrichments for each trend. Experimental design is shown on the bottom two rows, with the time points (i) post 0.05% deltamethrin WHO tube exposure and (ii) matched unexposed controls. Dark rings represent darkness in the 12:12 photoperiod.

Four of the 20 clusters showed sustained changes in the expression profiles after sublethal exposure to pyrethroids. These clusters include cluster 17 which represents transcripts with a trend for sustained up-regulation, whilst clusters 12, 5 and 6 show the converse trend (Additional File 1). Cluster 17 is enriched for DNA repair related transcripts (p=0.0084); whereas cluster 12 (which shows the clearest pattern of sustained down regulation) is enriched in mitochondrial electron transport chain (p=9.9E-4) consistent with oxidative damage and the associated reactive oxygen species burst shown to be caused by pyrethroid exposure in mammalian systems [34]. Clusters 5 and 6 are similarly enriched in respiratory related processes including carbon metabolism (p=3.6e-6; 3.2e-5), glycolysis (p=1.9e-5), oxidative phosphorylation (p=2e-12) and citrate (TCA) cycle (p=0.013; 3.7e-5) and indicative of an overall reduction in respiration post-insecticide exposure (Figure 1; Additional File 2).

Transcripts demonstrating consistent and significant up-regulation across all time points are listed in Table 1 and include the ortholog of the p53 transcription factor (AGAP002352-RB), which responds to genotoxic stress [35], AGAP001116-RA a D-amino acid oxidase linked with hydrogen peroxide production and detoxification, the UDP-transferase *UGT308G1* (AGAP007990-RA), *NET1* a nuclear associated protein (AGAP007948-RA) and the homolog of *galla-1* (AGAP007363-RA). Significant sustained down regulation across all time points was seen in 26 transcripts, including *CYP4H18*, the homolog of *Drosophila* transcription factor ticktock and the ionotropic receptor Ir7u (Additional File 3).

**Table 1:**
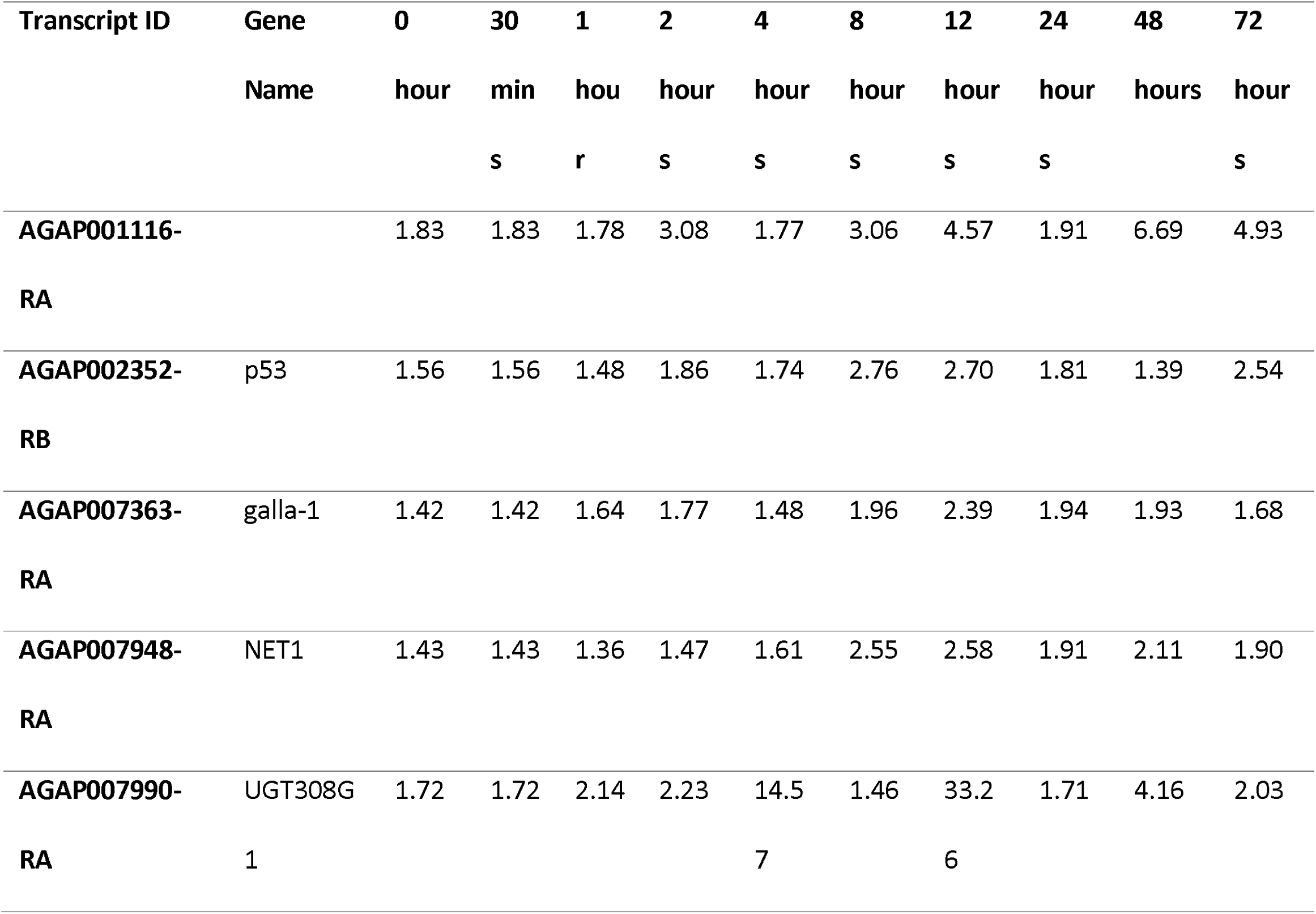
Transcripts significantly upregulated across all time points post-exposure. Transcript ID and fold changes compared to unexposed mosquitoes across each time point for transcripts with adjusted p < 0.05.

Four clusters grouped transcripts with clear changes in expression from 4-hours post-exposure (Additional File 1). Cluster 7 shows sharp down regulation 4 hours post-exposure and is highly enriched in immunoglobin-like proteins (p=2e-6), calcium ion binding (p=2.3e-5) and ion transport (p=0.0018); a number of these transcripts belong to a class of proteins called defective proboscis extension response. In *Drosophila* these proteins are neuronal and have been shown to be involved in responses to chemical stimuli and stress [36]. Similarly, 2 protein orthologs of *Drosophila* sidestep are represented in this cluster along with the interactor beat both of which have been linked to oxidative stress response induced locomoter defects [37]. Conversely, clusters 2 and 4 demonstrate strong induction of transcripts from 4 hours post-exposure and are enriched in translation (p=0.028) and structural components of the ribosome (p=1.5e-4) cluster 8 similarly shows induction from 4 hours post-exposure and is enriched in DNA replication (p=2.5e-4) indicating the onset of protein production related to insecticide exposure (Figure 1; Additional File 2).

Of those transcripts showing consistent significant expression directionality, 18 transcripts show a sustained, up-regulation from 1 hour or 2 hours post-exposure until 72 hours post exposure (Additional File 4), including the transcription factor *Dr* (AGAP003669-RA), which plays a role in locomotor activity and neuronal patterning in *Drosophila* and IMD a key immune-response regulator (AGAP004959-RB). A larger number of transcripts (265; 266 probes), show a delayed but sustained induction response beginning at 4- or 8-hours post exposure (Additional File 5). Of the 21 transcripts showing delayed, sustained down regulation from 1 or 2 hours onwards four are cuticular proteins (*CPLCG3, 4* and *15* and *CPR109*); this transcript list also includes the *D7r2* salivary protein (Additional File 4). A further 133 transcripts (140 probes) show sustained down-regulation either 4 or 8-hours post exposure (Additional File 5). Similarly, these transcripts contain a number of cuticular related transcripts including *CPR10, CPLCA1, CPLCX3, CPR59, CPCFC1* and *CPR132* and alternative probes for *CPCLG4*.

Two clusters show clear up-regulation of transcripts from 24-hours post-exposure (no clusters show a pattern of down-regulation after this time point although 43 transcripts show significant down-regulation from 24-hours onwards and 20 show significant up-regulation (Additional File 6)). Cluster 13 is enriched in oxidoreductase activity (p=0.0076), glutathione transferase activity (p = 0.027) and cytochrome p450 domains (p=0.016). Cluster 20 shows a similar expression pattern showing changes in transcripts related to lipid metabolism and biosynthesis (p=0.0028) (Figure 1); indicating that exposure to insecticide may lead to long term up-regulation of detoxification transcripts and differential expression of fatty acids (Figure 1; Additional File 2).

Two other clusters show strong enrichments but do not show a strong sustained temporal expression change. Clusters 16 and 19 show a peak of expression at 48- and 12-hours respectively (Additional File 1; Additional File 2). Cluster 19 is likely to represent strong circadian changes and is enriched in response to insecticide (p = 4.8e-2) and cytochrome p450s (p = 2.4e-6), likely reflecting the diel nature of expression of metabolic enzymes, described below. Cluster 16 is enriched in glutathione metabolic process, oxioreductase activity, cytochrome p450s and carboxylesterase (p = 7.11e-3; 1.4e-3; 7.2e-3; 1.5e-2) indicating changes relating to insecticide response peak strongly at 48-hours.

### Induction of gene families associated with pyrethroid resistance

Of 113 cytochrome p450s in the *Anopheles* genome, 81 are differentially expressed in at least one timepoint post pyrethroid exposure (Supplementary Table 6). Of the 8 cytochrome p450s that bind to pyrethroid insecticides and have been widely implicated in pyrethroid resistance [16, 38] (Figure 2), two, (*CYP6M2* and *CYP6Z2*), are strongly induced after deltamethrin exposure. Closely related P450s not previously associated with pyrethroid metabolism are also strongly induced (*CYP6M1, CYP6M3, CYP6Z3*) (Supplementary Table 6). Several other p450s are induced over multiple hours or days, including *CYP4G16* and *CYP4G17*, both linked with cuticular thickening [12], *CYP4D17, CYP6AH1, CYP6Z1* and *CYP4C27*. Notable genes from other detoxification gene families that are induced post exposure include, *GSTD1, ABCG5* (previously shown to enriched in the abdomen and up-regulated across multiple resistant population [18]), *ABCC14* (the homolog of *Drosophila* multidrug resistance protein 1 and up-regulated in multiple resistant populations [18]), *COE13O* and *UGT308G1*. CSPs have recently been linked with pyrethroid resistance in West Africa [13]; *SAP3*, is highly induced from 8 hours (Additional File 7) whereas SAP2 is significantly over expressed at a single timepoint (8 hours).

**Figure 2:**
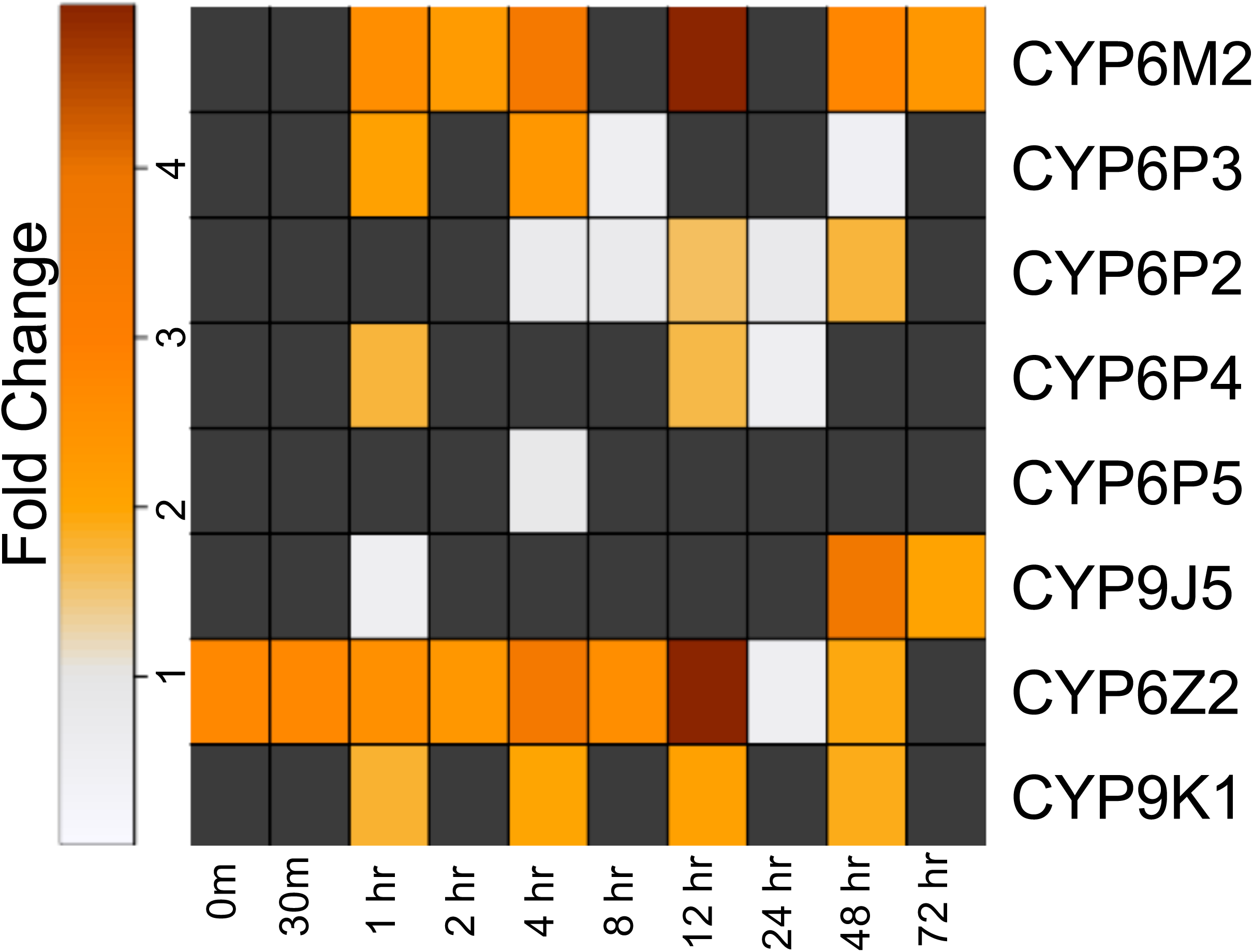
Cytochrome p450 pyrethroid metabolisers. Transcript expression level for 8 cytochrome p450s that have previously been shown to bind insecticide directly [16, 38, 39]. Dark grey boxes represent non-significant transcripts.

Trends of down-regulation are also seen within the detoxification families, including *CYP4H18* which was down-regulated across all time points. Other transcripts showing down regulation across multiple time points include two transcripts labelled as *CYP9M1* (AGAP009374-RA and AGAP009363-RA), *GSTZ1* (-RA only), *GSTD12, GSTE1, GSTE2, GSTMS3, GSTS1, ABCC7, COEAE5G* and *UGT301E2*. Interestingly, *GSTE2* has been strongly linked with DDT and pyrethroid resistance [17]; however, in this strain it is strongly down regulated from 4 to 48 hours post-exposure.

### Changes to respiratory-related transcripts

As shown using soft clustering, transcripts demonstrating sustained downregulation post-pyrethroid exposure are enriched in transcripts involved in both the mitochondrial oxidative phosphorylation chain and in the TCA cycle, indicating a wider change to respiration caused by pyrethroid exposure. The changes in gene expression are shown in Additional File 8 and Figure 3 provides a pictorial representation of the oxidative phosphorylation chain from which it can be seen that pyrethroid exposure suppresses gene expression for each of the five members of the respiratory complex (Figure 3).

**Figure 3:**
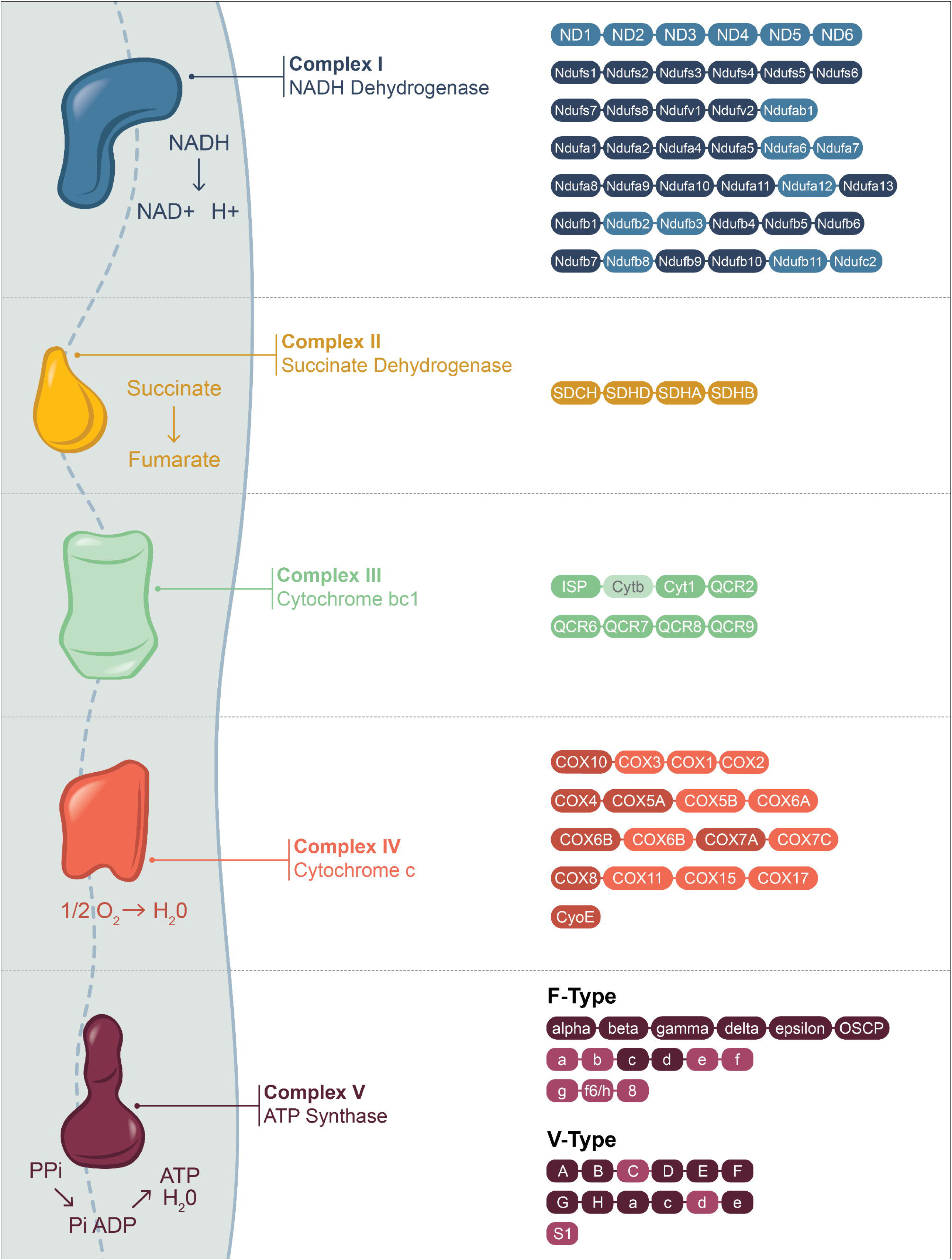
Transcripts in the oxidative phosphorylation pathway down regulated by pyrethroid exposure. Modified KEGG pathway showing all transcripts in the oxidative phosphorylation in *An. gambiae* (KEGG organism: aga). Darkened boxes represent transcripts that are significantly down regulated in at least one time point in the time course data. ND1-6 are not represented on the microarray as they are mitochondrial genes.

### (ii) Ageing increases susceptibility to insecticides due to changes in expression of insecticide related transcripts

A total of 931 transcripts (1033 probes; representing 6.93% of the array) were significantly differentially regulated between the 0-hour (3 day old) and 48-hour (5 day old) timepoints in unexposed mosquitoes, indicating the extensive changes in gene expression as female mosquitoes age (Additional File 9). Of these, 403 transcripts (449 probes) were up-regulated and 528 (584 probes) down-regulated. Both up and down regulated genes showed significant enrichment for GO terms related to detoxification. Genes involved in heme-binding (p = 0.0085), glutathione transferase (p = 0.0011) and glutathione peroxidase activity (p= 0.049) were up-regulated whereas downregulated genes were enriched in insecticide catabolic process (p = 0.000494), mono-oxygenase activity (p = 3.8e-6), iron-binding (p = 2.2e-7), oxidoreductase activity (p = 1.5e-5), heme-binding (1.5e-5) and organic anion transporter (p = 2.6e-4). A number of detoxification genes (274 transcripts, representing < 2% of the genome) (Table 2), chemosensory proteins [13] and a cuticular protein [40] previously linked with insecticide resistance are expressed a lower levels in 5 day females than 3 day olds, perhaps providing an explanation for previous observations that resistance to pyrethroid insecticide falls with mosquito age [41, 42]. However, some resistance-related transcripts, including methoprene tolerant [21], several members of the GSTD and GSTE families and CSP6 [13] are up-regulated in older mosquitoes.

**Table 2:** Cytochrome p450s down regulated in 5 day vs 3 day old females. Transcript ID, Gene Name, Adjusted p value and absolute Fold Change of cytochrome p450s previously implicated in insecticide resistance [14–16, 21, 43] in 5-day old adult female mosquitoes compared to 3-day old. Asterisk’s indicates pyrethroid metabolising enzymes [14–16].

| Transcript ID | Gene Name | Adjusted p Value | Fold Change |
| --- | --- | --- | --- |
| AGAP000088-RA | CYP4H19 | 0.012 | 0.294 |
| AGAP000818-RA | CYP9K1* | 0.013 | 0.447 |
| AGAP000877-RA | CYP4G17 | 0.042 | 0.556 |
| AGAP001076-RA | CYP4G16 | 0.036 | 0.634 |
| AGAP001076-RB | CYP4G16 | 0.050 | 0.646 |
| AGAP001076-RC | CYP4G16 | 0.045 | 0.721 |
| AGAP002862-RA | CYP6AA1 | 0.049 | 0.590 |
| AGAP002865-RA | CYP6P3* | 0.006 | 0.379 |
| AGAP002894-RA | CYP6Z4 | 0.041 | 0.747 |
| AGAP008212-RA | CYP6M2* | 0.004 | 0.382 |
| AGAP008213-RA | CYP6M3 | 0.009 | 0.451 |
| AGAP008214-RA | CYP6M4 | 0.043 | 0.739 |
| AGAP008217-RA | CYP6Z3 | 0.042 | 0.524 |
| AGAP008218-RA | CYP6Z2* | 0.029 | 0.373 |
| AGAP008219-RA | CYP6Z1 | 0.008 | 0.336 |
| AGAP013490-RA | CYP4H24 | 0.013 | 0.275 |

### iii) Diel cycle controls the expression of insecticide resistance transcripts

*Anopheles* mosquitoes are night biting mosquitoes and, as adults, are most likely to encounter insecticide when searching for a blood meal in the evening. To assess the diel nature of insecticide related transcripts in a multi-insecticide resistant population, age matched females were compared in two steps (i) comparing transcript expression in mosqutioes sacrificed at 7pm or 11pm (8hrs and 12hrs) and (ii) 11pm or 11am (12hrs and 24hrs) (Figure 1). No transcripts showed a significantly differential expression pattern in group (i); however, in group (ii) 506 (587 probes) transcripts showed differential expression (Additional File 10). Of these 230 (273 probes) were up-regulated and 276 (314 probes) were down-regulated in the morning compared to the evening. Transcripts overexpressed in the morning were enriched in oxidoreductase and monooxygenase activity (p = 2.6e-5; 3.5e-5); heme binding (p = 3.9e-5); iron ion biding (p = 2.04e-4) and both glutathione peroxidase and transferase activity (p = 3.54e-4; p = 8.1e-4). Transcripts overexpressed in the evening were similarly enriched in oxidoreductase and monoxygenase activity (p = 6e-4; 9.3e-4), heme binding (p = 6.5e-4), iron ion binding (p = 5.4e-4) and insecticide catabolic process (p = 0.0014). Within these transcripts are direct insecticide interactors including *SAP2* [13], *CYP6M2* [15, 16], *CYP6P3* [14, 16], *CYP6P4* [16], *CYP6Z1* [43] and *CYP9K1* [39] (Table 3), all of which are expressed at higher levels at 11pm than 11 am. As detoxification-related transcripts are enriched both in the morning and in the evening, this could indicate a two-phase process in metabolic clearance of pyrethroid insecticides with a subset of cytochrome p450s catalysing the initial oxidation reaction being highly enriched at night (p = 3.05e-7; 0.013), and a separate subset of cytochrome p450s, plus COEs and GSTs, responsible for secondary pyrethroid metabolism enriched in the morning (p = 4.37e-10; 5.2e-3; 2.56e-8). Further, Anopheline antiplatelet protein, four salivary gland related proteins and several trypsin transcripts are enriched at night-time when the mosquito would be seeking a bloodmeal. *Cycle* and *Clock* are upregulated whilst *Period, Cryptochrome 2, Timeless* and *PDP1* are downregulated in this dataset, confirming the rhythmic nature of these changes following the pattern previously reported in *An. gambiae* [44].

**Table 3:**
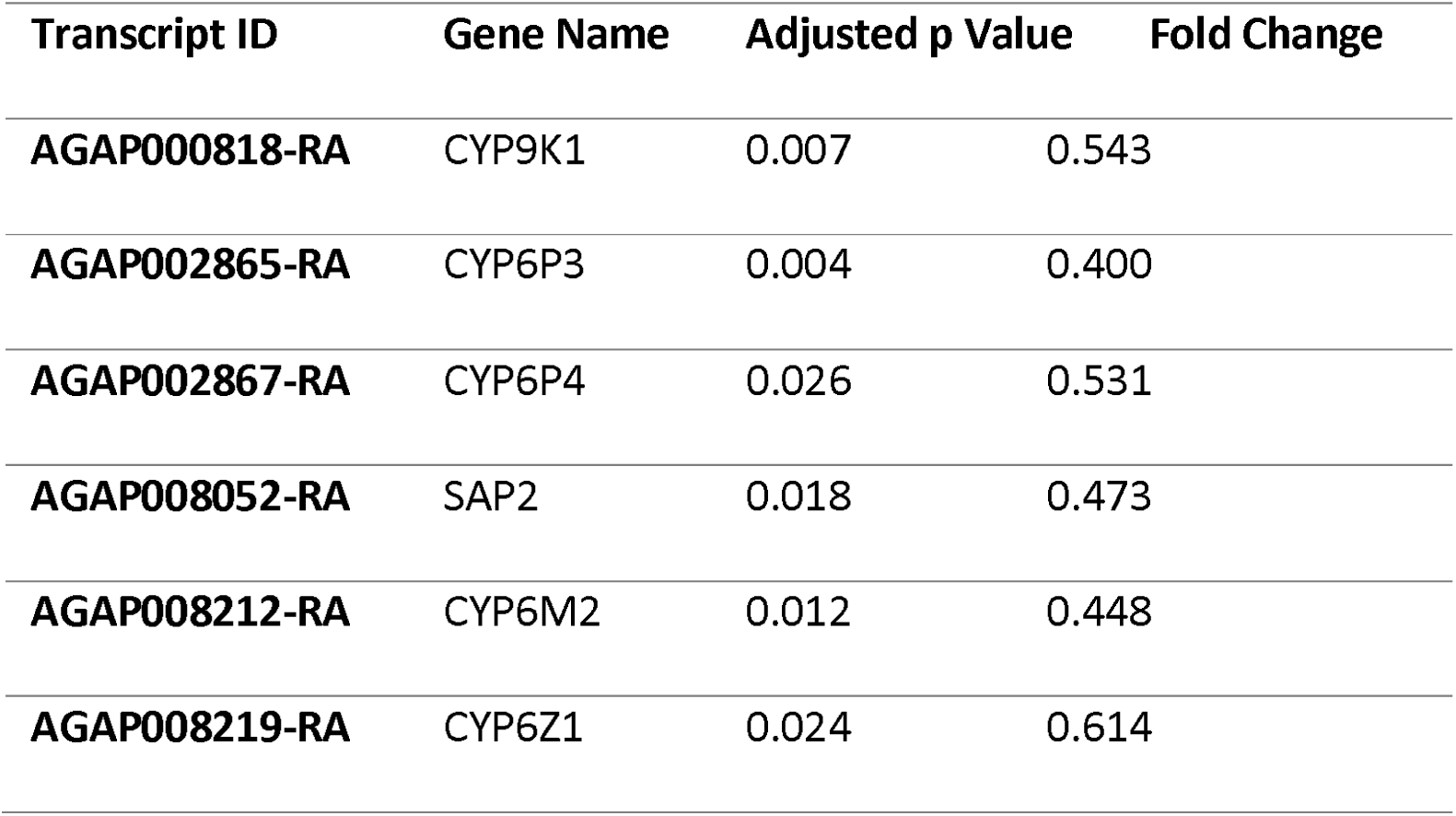
Direct pyrethroid interactors upregulated at night. Transcript ID, Gene Name, Adjusted p value and absolute Fold Change of transcripts that directly interact with insecticides showing enrichment at 11pm compared to 11am (downregulation at 11am compared to 11pm as shown above).

## Discussion

Insecticide resistance has been defined by WHO as the number one obstacle to malaria elimination. The majority of studies investigating the molecular basis of resistance focus on the constitutive overexpression of transcripts in resistant mosquito populations, compared to susceptible controls; however, induction of gene expression after a sub-lethal dose of insecticide is likely to be equally important for long term fitness effects and parasite transmission. In this study, we explore these factors using whole transcriptome microarrays with a pyrethroid-resistant *An. coluzzii* population, VK7. This strain was selected as the high levels of pyrethroid resistance are conferred by multiple mechanisms including target site mutations and high levels of overexpression of the cytochrome p450 pyrethroid metabolisers *CYP6M2* and *CYP6P3* [33] with further, less well characterised resistance mechanisms potentially associated with increased expression of α-crystallins and an F-Type ATPase [33].

In this study we identify five phases to the response to pyrethroid exposure. There is an immediate and sustained reduction in genes involved in mitochondrial respiration and a sustained increase in expression of DNA-damage response related transcripts. A reduced respiratory rate after exposure to pyrethroid insecticides has been widely reported in mammalian and fish systems through inhibition of the oxidative phosphorylation chain [45–47]; however, as far as we are aware, no studies have examined this in insects. It is possible a reduction in respiration post-pyrethroid exposure represents a compensatory mechanism to reduce mitochondrially produced reactive oxygen species due to exogenous ROS production from sub-lethal pyrethroid exposure [48]. Similarly, the oxidative stress caused by pyrethroid exposure are likely to cause genotoxicity [48, 49], hence explaining the up-regulation of DNA-repair related transcripts. We further show that 4-hours post exposure there is a large change in transcription with up-regulation of translation and down regulation of ion transport and immunoglobin-like proteins. The up-regulation of transcripts related to translation could be due to the sustained changes seen in transcriptional activity for up to 72 hours resulting in higher levels of protein production. Further, the down-regulation of neuronal-related transcripts could account for some behavioural changes seen in *Anopheles* mosquitoes on contact with ITNs [50, 51]. Perhaps most surprisingly, enrichment in monooxygenase activity, cytochrome P450 domains and glutathione activity, commonly associated with insecticide resistance are only induced after 24-hours post exposure. The induction of these transcripts from a day post-exposure across all subsequent time points suggests that sub-lethal exposure could lead to higher levels of resistance upon second exposure the following night through overactivity of detoxification related transcripts. Indeed, this has been demonstrated after mosquitoes take a bloodmeal [52], which induces a large oxidative stress response, similar to those seen in mammalian systems after pyrethroid exposure.

Five transcripts show differential expression from immediately after pyrethroid exposure to the maximal timepoint, 72 hours. These transcripts are likely to be some of the most important for insecticide response and contain p53, a DNA-damage related transcription factor [35], *UGT308G1*, the homolog of *galla-1, NET1* and a transcript linked with reactive oxygen species response, a D-amino-acid oxidase [53]. *p53* has been shown to have multiple roles in cellular response to genotoxic stress in *Drosophila* [35]. Few studies have explored the function of these genes in *Anopheles* mosquitoes. One study on mosquito p53 orthologs describes a direct role in response to oxidative stress upon arboviral infection [54], perhaps indicating that this gene may respond to similar stress post-pyrethroid exposure. Similarly, the UGT family have previously been linked to insecticide resistance [20], the high level of induction of this transcript across all timepoints suggests that the role of this family in pyrethroid detoxification merits further study. Indeed, the UGT308 family has been shown to be an essential family for the biotransformation of pyrethroid insecticides in *An. sinensis*, an Asian malaria vector [55]. A SNP in *galla-1* was found to be significantly associated with *Drosophila* response to oxidative stress [37] and was also found to play a role in protective response to reactive oxygen species in fragile genetic sites through alterations to aerobic metabolism [56]. *NET1* has a GO Term of protein binding and is found in the KEGG pathway for ribosome biogenesis, indicating a role in translation. The *Drosophila* homolog of the D-amino-acid oxidase described here is localised to the peroxisome, these membrane bound organelles play a key role in both the production and detoxification of cellular reactive oxygen species [53]. Taken together, there is strong indication that these transcripts play a key role in response to oxidative stress caused by exposure to pyrethroid insecticides, either through maintaining cellular homeostasis or protection of genetic material. The down-regulated transcripts include a number of cuticular proteins and the ABC transporter *ABCH2*, a half transporter whose role in insects is poorly characterised [18]. Interestingly, the salivary protein *D7r2* has previously been linked to bendiocarb resistance through ubiquitous overexpression [57]; however, these data indicate that this transcript may not be important in pyrethroid resistance in this population, supported by recently published data from *An. gambiae* in Cameroon [58].

The induction of detoxification candidates previously shown to be involved with insecticide resistance is an important consideration, as many of these transcripts are already expressed at constitutively higher levels within pyrethroid resistant mosquito populations. For example, *CYP6M2* is 8 fold constitutively overexpressed in VK7 compared to a susceptible control [33], and here we show a further 5.2 fold overexpressed maximally post-exposure. Similarly, *CYP6Z2* is 3.5x constitutively overexpressed [59] and a further 7.2x maximally induced, clearly demonstrating the importance of induction of these pyrethroid metabolisers.

Pyrethroid resistance has previously been shown to fall with age [41, 42], in the absence of a blood meal [52]. To explore the transcriptional basis of this reduction, we compared 3- and 5-day old mosquitoes and found substantial changes in transcript expression. Of the transcripts down regulated were a number of genes previously linked to insecticide resistance such as, a cuticular protein *CPLCG5* [40], *CYP6P3* [14], *CYP4G16* [12] and the chemosensory protein *SAP2* [13]. Given the relatively large reductions in transcript expression over a short time period, it is likely that the reduction in expression of these key transcripts play a large role in the relative loss of resistance; however, further time points, with accompanying bioassays, would be needed to investigate further. Previous data has demonstrated a high number of transcripts are controlled by the diel cycle in susceptible *An. gambiae* mosquitoes, with an enrichment of those involved in detoxification [44]; here we show that *CYP6Z1, CYP6P3* and *CYP6M2* showed peak expression during the evening [44] and *SAP2*, a key pyrethroid resistance determinant in West Africa, is also upregulated at night. There is also a clear enrichment for oxidoreductase activity early morning, which may indicate a potential two-step process in metabolic clearance of insecticides, with the first being direct metabolism through cytochrome p450s and binding and the second stage related to clearance of the metabolites through up-regulation of GSTs.

Although this study provides a comprehensive picture of sub-lethal pyrethroid exposure in a resistant *An. coluzzii* population from Burkina Faso, it is clear that even within country there are differing resistance mechanisms between sites [33]. Further, the 1-hour WHO exposure used here is not representative of pyrethroid dose received under natural settings, where mosquitoes spend shorter time in contact with higher concentrations on insecticide treated surfaces [50].

## Conclusion

This study provides insight into the large changes in transcript expression at various time-points post-pyrethroid exposure. The sustained transcriptional changes seen here are likely to have important phenotypic effects; for example, the reduced respiratory rate, and increased investment in DNA repair and protein production are likely to have energetic costs which may reduce the fitness of mosquitoes post pyrethroid exposure. Although no shortening of longevity is seen in this population after insecticide exposure in a laboratory environment [6], other fitness traits such as fertility and fecundity merit further studies. Further, if pyrethroid exposure impacts the redox state of the mosquito, as indicated by the widespread changes to oxidoreductase-related transcripts and respiratory rate, this may impact the mosquito’s ability to transmit pathogens. Disruption of parasite development due to changes in redox state has been shown experimentally through reducing catalase activity which in turn reduces oocyst density in the midgut [60], whilst the initial immune response to parasite invasion is a large reactive oxygen species burst [60, 61]. Hence, future studies should investigate how pyrethroid exposure affect the development of pathogens in the mosquito.

## Methods

### Mosquito Rearing Conditions

The *An. coluzzii* used in these experiments were all presumed mated and reared under standard insectary conditions at 27 °C, 70-80% humidity and 12:12 hour photoperiod. The VK7 colony was originally collected from Vallee de Kou, Burkina Faso and has been maintained under pyrethroid selection pressure at Liverpool School of Tropical Medicine since 2014 [33, 59]. Resistance in this population is regularly characterised and demonstrates high levels of pyrethroid and DDT resistance [33].

### Insecticide Exposures

Pools of 20-30 3-day old adult females from the same generation were exposed for 1-hour to 0.05% deltamethrin impregnated papers in a WHO tube bioassay as previously described [62]. Mosquitoes used to look at diel cycle and ageing were taken from a different generation to exposed mosquitoes due to high mosquito numbers needed. The starting point for all assays was 10am.

### Microarray Experiments

RNA was extracted from pools of 7-10 adult females from unexposed VK7 at the following time points: 3 day old (‘0 hour’); 8-hours, 12-hours, 24-hours, 48-hours and 72-hours and exposed VK7 at the following time points: immediately after 1 hour exposure and then 30 minutes, 1 hour, 2 hours, 4 hours, 8 hours, 12 hours;,24 hours and 48 hours post exposure. Each biological replicate consisted of pooled RNA, extracted using PicoPure RNA Isolation kit (Arcturus) following manufacturer’s instructions. Three or four biological replicates were prepared for each time point for each strain. All biological replicates were taken from the same colony cage; for each time point, all replicates were from mosquitoes that had been exposed to deltamethrin simultaneously. Each timepoint for the exposed mosquitoes was competitively hybridised with the previous time point. The unexposed mosquitoes were competitively hybridised as follows: 0-hours vs 48-hour (ageing); 8-hour vs 12-hour and 12-hour vs 24-hour (two diel time points). The quality of the RNA was assessed using a nanodrop spectrophotometer (Nanodrop Technologies UK) and TapeStation (Agilent). 100ng of RNA was amplified and labelled with either *Cy3* and *Cy5*, using the ‘Two color low input Quick Amp labelling kit’ (Agilent) following manufacturer’s instructions. Samples were then purified using the RNA purification kit (Qiagen), with cRNA yield and quality assessed using the spectrophotometer (Nanodrop Technologies UK) and TapeStation (Agilent). Microarray data was obtained from scanning 15k Agilent *Anopheles* microarrays (ArrayExpress accession number A-MEXP-2196), hybridised with labelled cRNA, with an Agilent G2205B scanner. Hybridisations were carried out over 17 hours at 65° C at 10rpm rotation and washed following manufacturer’s instructions (Agilent). Datasets are available at ArrayExpress: exposure time course (E-MTAB-9422) and ageing time course (E-MTAB-9423).

### Data Analysis

The resulting microarray data was analysed by fitting linear models to normalised corrected signals using the R package limma [63] following package instructions. Briefly, within and between array normalisation was carried out using loess and Aquantile respectively, with background correction using normexp. Microarrays were analysed both as hybridised using lmFit and eBayes for the unexposed mosquitoes, and also using separate channel analysis for two-colour data allowing comparison to the unexposed control as described in package instructions using intraspotCorrelation, lmscFit and eBayes (Github: https://github.com/VictoriaIngham/Time_Course). Benjamini and Hochberg adjusted p value of less than or equal to 0.05 was used for probe significance. Clusters of probes following the same expression patterns were found using Mfuzz [64] with 20 clusters. Mfuzz was selected over k-means clustering as it utilises soft clustering, allowing a membership score to be assigned to each transcript. Optimal fuzzifier value was calculated following published guidelines [65] (m = 1.30), optimal clusters calculated using sum of squares error (c = 7-10); c = 20 was used to ensure all substructure identified and no between group correlation > 0.9. All data was standardised so that the expression profile for each transcript had a mean on 0 and a standard deviation of one. All enrichment analyses were carried out using DAVID [66] and KEGG [67] using Benjamini and Hochberg adjusted p value of less than or equal to 0.05 for significance. *Drosophila* homologs were found using VectorBase [68] and FlyBase [69]. Heatmaps were produced using ggplot2, in the case of multiple probes for individual transcripts, these were averaged across all probes. In these cases, significance for multiple probes was defined as at least one of the four probes show significance.

## Abbreviations

ITNs: Insecticide Treated Bed Nets
WHO: World Health Organisation
Kdr: Knockdown Resistance
CSPs: Chemosensory Proteins
P450s: Cytochrome P450s
GSTs: Glutathione-s-transferases
ABCs: ABC transporters
COEs: carboxylesterase
UGTs: UDP-glucuronyl transferases
TCA: Citrate Cycle
KEGG: Kyoto Encyclopaedia of Genes and Genomes
GO: Gene Ontology
SNP: Single Nucleotide Polymorphism

## Declarations

### Ethics approval and consent to participate

Not Applicable

### Consent for publication

Not Applicable

### Availability of Data and Materials

The datasets supporting the conclusions of this article are available in the ArrayExpress repository: exposure time course (E-MTAB-9422) and ageing time course (E-MTAB-9423). All R code used in this analysis is available from the corresponding author upon request and at https://github.com/VictoriaIngham/Time_Course.

### Author Contributions

VAI and HR conceived the study and drafted the manuscript. FB performed all mosquito maintenance, insecticide exposures and RNA extractions. VAI performed the microarray experiments and all subsequent analysis. All authors have read and approved the manuscript.

## Acknowledgements

We thank Manuela Bernardi (LSTM) for preparation of Figures 1 and 3 and John Morgan (LSTM) and Kobie Hyacinthe Toé (Centre National de Recherche et de Formation sur le Paludisme) for original mosquito collections that formed the VK72014 colony.

---

**Additional File 1: Mfuzz clusters for all transcripts**. Expression patterns for all transcripts across the 20 soft Mfuzz clusters. Red indicated high cluster membership, blue intermediate and green low. Y-axis indicated normalised expression change and the x-axis represents each time point.

**Additional File 2: Mfuzz cluster membership and associated enrichments**. Cluster ID corresponding to visual display, significant GO term enrichments for each of Biological Processes, Cellular Component and Molecular Function, significant KEGG pathway enrichment and significant InerPro domains. Pattern describes the general pattern seen in the cluster. All p values are shown in brackets with BH adjustment.

**Additional File 3: Transcripts significantly down regulated at all time points**. Transcript ID, gene name, gene description, fold change and adjusted p value for each time point.

**Additional File 4: Transcripts significantly differential from 1- or 2-hours post-exposure onwards**. Transcript ID, gene name, gene description, fold change and adjusted p value for each time point. Red indicates non-significance.

**Additional File 5: Transcripts significantly differential from 4- or 8-hours post-exposure onwards**. Transcript ID, gene name, gene description, fold change and adjusted p value for each time point. Red indicates non-significance.

**Additional File 6: Transcripts significantly differential from 24 hours post-exposure onwards**. Transcript ID, gene name, gene description, fold change and adjusted p value for each time point. Red indicates non-significance.

**Additional File 7: Changed in pyrethroid resistance-related gene families**. Transcript ID, Gene Name and Log2 fold change across all time points for each transcript in the six resistance-related families that are significantly (adjusted p<= 0.05) differential in at least one time point. Black boxes represent non-significant time points. Transcript ID -RX accounts for different splice variants. Heatmap with colour key representing raw fold change for each transcript, in the case of multiple probes the average fold change across all probes. Tabs on the excel sheet represent different families.

**Additional File 8: Significant changes in respiratory-related transcripts**. Heatmaps showing transcripts involved in (A) Oxidative phosphorylation and (B) TCA cycle that are differential in at least one time point. Pathway membership as defined by KEGG. Transcript ID followed by generic name is shown in row labelling, columns represent different timepoints. Dark grey indicated non-significant.

**Additional File 9: Significantly differentially expressed transcripts between 3- and 5-day old females**. Transcript ID, Gene Name and Gene Description (VectorBase, April 2020), adjusted p-value and fold change for each transcript. Down-regulation indicates transcripts with lower expression in older females, up-regulation is the converse.

**Additional File 10: Significantly differentially expressed transcripts between night-time (11pm) and morning (11am)**. Transcript ID, Gene Name and Gene Description (VectorBase, April 2020), adjusted p-value and fold change for each transcript. Down-regulation indicates transcripts with higher expression in the evening, up-regulation is the converse.

