## Supplementary figures and images for "Sub-lethal pyrethroid exposure and ageing lead to pronounced changes in gene expression in insecticide resistance *Anopheles coluzzii*"

### Additional File 8.png

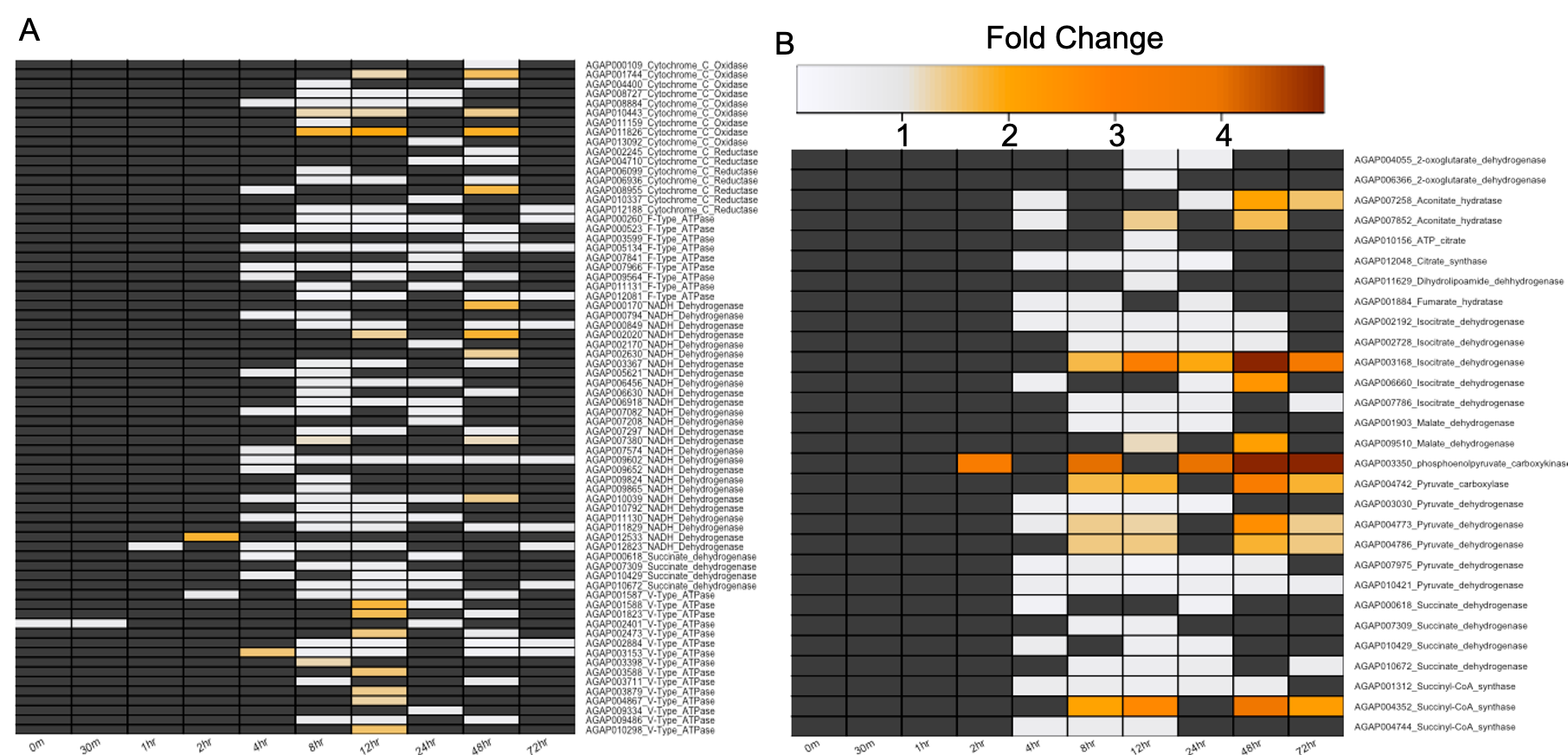
